# Reward-driven cerebellar climbing fiber activity influences both neural and behavioral learning

**DOI:** 10.1101/2024.10.09.617466

**Authors:** Shuyang Jin, Court Hull

## Abstract

The cerebellum plays a key role in motor coordination and learning. In contrast with classical supervised learning models, recent work has revealed that CFs can signal reward-predictive information in some behaviors. This raises the question of whether CFs may also operate according to the principles of reinforcement learning. To test how CFs operate during reward-guided behavior, and evaluate the role of reward-related CF activity in learning, we have measured CF responses in Purkinje cells of the lateral cerebellum during a Pavlovian task using 2-photon calcium imaging. Specifically, we have performed multi-stimulus experiments to determine whether CF activity meets the requirements of a reward prediction error (rPE) signal for transfer from an unexpected reward to a reward-predictive cue. We find that once CF activity is transferred to a conditioned stimulus, and there is no longer a response to reward, CFs cannot generate learned responses to a second conditioned stimulus that carries the same reward prediction. In addition, by expressing the inhibitory opsin GtACR2 in neurons of the inferior olive, and optically inhibiting these neurons across behavioral training at the time of unexpected reward, we find that the transfer of CF signals to the conditioned stimulus is impaired. Moreover, this optogenetic inhibition also impairs learning, resulting in a deficit in anticipatory lick timing. Together, these results indicate that CF signals can exhibit several characteristics in common with the rPEs that have been observed during reinforcement learning, and that the cerebellum can harness these reward-related learning signals to generate accurately timed motor behavior.

## Introduction

Climbing fibers (CFs) play a key role in many forms of cerebellar learning, and are thought to provide instructional signals that generate synaptic plasticity and modify behavior according to the principles of supervised learning^1^. Recent work has revealed that CFs can also display characteristics that are typically associated with reward-prediction errors (rPEs), the instructional signal necessary for reinforcement learning, in some behaviors guided by predictions about reward^2,3^. In particular, it has become clear that CFs can respond to unexpected reward, and generate learned responses to reward-predictive cues^4–8^. However, CFs also exhibit properties that are inconsistent with leading models of reinforcement learning, such as elevated spiking in response to omitted reward^7^. Thus, while many properties of CF signaling in reward tasks are consistent with rPEs, they may also be consistent with signaling other information such as stimulus salience, valence or novelty.

To determine how CFs generate teaching signals in reward-based tasks, and what similarities these signals have with classically described rPEs, it is important to test whether they obey key principles of that have been described in other brain structures such the ventral tegmental area (VTA) thought to be necessary for reinforcement learning. There, some dopamine (DA) neurons are thought report rPEs in reward-guided tasks^9^, and to follow many of the formal predictions established by reinforcement learning theory^10–12^. For example, DA neurons can signal positive rPEs (increased spiking) for unexpected outcomes that are better than anticipated (surprise reward delivery), and these responses shift with learning to represent sensory stimuli that predict such outcomes^13^. In accordance with the formalism of leading reinforcement learning theories such as temporal-difference learning (TD learning), this shift requires the initial response to unexpected reward. In the VTA, this has been demonstrated using so-called ’stimulus blocking’ conditions that leverage the absence of reward-driven spiking in response to a fully trained sensory cue^14^. Because reward-driven spiking is abolished in this condition, a new reward-predictive stimulus that carries the same prediction cannot generate learned DA neuron responses.

To assess the similarity of cerebellar CF responses to those of DA neurons, and test whether CFs follow these key principles of reinforcement learning, we have used a stimulus-blocking paradigm to test the requirements for generating learned CF responses to reward-predictive sensory cues. We find that, like canonical DA neurons of the VTA, cerebellar CFs do not learn sensory stimuli that carry fully redundant reward predictions. To test the causality of reward-driven CF responses in generating learned, reward-predictive sensory responses, we have optogenetically inhibited CF activity across learning, preventing the initial response to unexpected reward. These experiments impair the transfer of CF responses to a reward-predictive auditory stimulus, suggesting a potential role of cerebellar learning in generating predictive CF activity. Finally, we also observe an impairment of learned lick-timing after inhibiting reward-driven CF activity, suggesting a role for these teaching signals in generating predictively timed movements. Together, these results reveal important links between cerebellar CF responses and the predictions of reinforcement learning theory, and indicate a key role for reward-driven CF responses in cerebellar learning.

## Results

To test the properties of climbing fiber activity during reward conditioning, we used a Pavlovian conditioning task optimized for Ca^2+^ imaging in the lateral cerebellar cortex^4^. Previously work indicated that mice can effectively associate a visual cue with reward^4^, and that CF activity is transferred to the visual cue with learning. To test whether CFs can also respond to other types of reward-predictive sensory stimuli, we trained animals on a version of the same task using an auditory conditioned stimulus (CS). Specifically, mice were trained to associate a tone (200 ms, 5 kHz, 75 dB) with a delayed saccharine water reward (unconditioned stimulus-US; 700 ms delay, **Figure 1A**).

**Figure 1.**
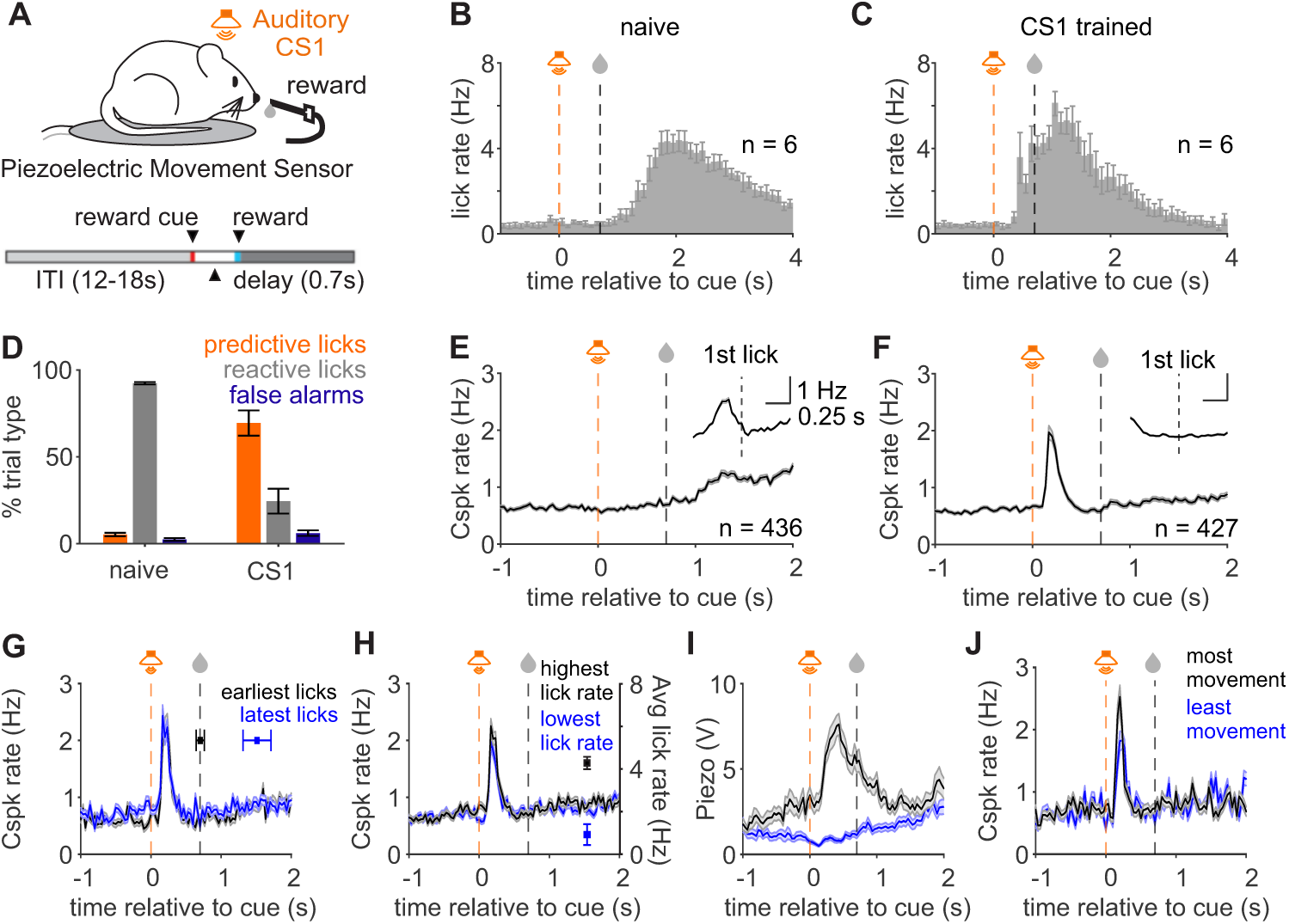
Pavlovian task to associate reward with a predictive auditory cue (CS1). **A)** Schematic of experimental design. Mice were presented with an auditory tone (200 ms, 5 kHz, 75 dB) prior to reward (700 ms after tone onset). **B,C)** Summary histograms of mean lick rates across experiments in naive (**B**) and trained (**C**) mice (n=6 mice). Dashed lines represent the time of cue and reward respectively. **D)** Summary of the fraction of trials across mice with predictive licks (orange), reactive licks (gray), or early licks (false alarms, blue) during the trial. **E)** Peristimulus time histogram (PSTH) of complex spike (Cspk) rate aligned to cue onset in naive animals (n=436 dendrites). Inset, Cspk rate aligned to the first lick after cue presentation. **F)** Same as E) but for trained mice (n=427 dendrites). **G)** PSTH of Cspk rates in trained mice for trials with the earliest and latest licks. Dots represent mean onsets for earliest (black) and latest (blue) 20% of lick bursts. **H)** Same as G) for trials with the highest (black) and lowest (blue) 20% of pre-reward lick rates. Dots represent mean lick rates for highest (black) and lowest (blue) 20% of lick rates. **I)** PSTHs of mean piezo voltage traces reflecting the 20% of trials with the most (black) and least (blue) animal movement **J)** Same as H) for Cspk rate on trials with the most (black) and least (blue) animal movement. All data are presented as mean +/- SEM.

Mice learned to associate the auditory cue with reward, developing robust licking to the auditory CS (**Figure 1B,C**) (training = 1089.3 ± 103.6 trials, 9 ± 0.7 sessions, n = 6 mice). To assess learning, we measured the fraction of trials with predictive and reactive licking, respectively defined as licking onset occurring before or after reward delivery. Learning resulted in a significant increase in predictive licking (Pre-learning: 5.2 ± 0.1%; Post-learning: 69.5 ± 7.3%; p = 3.52 × 10^-4^, n = 6 mice, paired t-test **Figure 1D, See also Figure S1**), and a decrease in reactive licking (Pre-learning: 92.4 ± 0.7%; Post-learning 24.5 ± 7.1%; p = 2.36 × 10^-4^, n = 6 mice, paired t-test, **Figure 1D, See also Figure S1**). In addition, mice exhibited significantly shorter reaction times (Pre-learning, 1491.7 ± 75.6 ms; Post-learning, 601.8 ± 78.3 ms; p = 8.16 × 10^-4^, n = 6 mice, paired t-test, **Figure S1**) and lower miss rates after learning (Pre-learning, 29.0 ± 6.7%; Post-learning, 4.0 ± 1.8%; p = 0.02, n = 6 mice, paired t-test, **Figure S1**).

Using this task, we measured complex spikes (Cspk) in the dendrites of Purkinje cells, as a proxy for CF activity^7^, in lobule simplex before and after learning (**Figure 1E,F, See also Figure S1**). Consistent with previous work^4^, we observed a significant Cspk response to unexpected reward delivery in naive animals, best revealed in the lick-triggered average of Cspk activity (p = 1.19 × 10^-7^, paired t-test, **Figure 1E, See also Figure S1**). In contrast, we did not observe a Cspk response to the auditory CS in naive animals (n = 436 dendrites, 6 mice, p = 0.12, paired t-test, **Figure 1E, See also Figure S1**).

After learning, a significant Cspk response occurred within 250 ms after the auditory CS (n = 427 dendrites, 6 mice, p = 5.51 × 10^-49^, paired t-test, one-tailed; time from cue onset to peak Cspk rate: 224.43 ± 2.31 ms, **Figure 1F, See also Fig S1**). In addition, Cspks no longer occurred following the first lick after reward (p = 0.08, paired t-test, **Figure 1F, See also Figure S1**). Because the lick-related Cspk response is absent after learning, the response in naive animals is likely not exclusively motor-related, and may involve reward anticipation or detection.

To further test the role of motor output in Cspk responses after learning, we segregated trials into subsets according to lick times and lick rates. Specifically, we averaged trials with the earliest and latest quartiles of lick onsets, and trials with the highest and lowest quartiles of lick rates (**Figure 1G,H**). This analysis revealed that the Cspk response had the same amplitude (p = 0.21, n = 427 dendrites, 6 mice, paired t-test, **Figure 1G**) and timing (time from cue on earliest lick trials = 212.41 ± 2.66 ms; latest lick trials: 208.27 ± 2.54 ms, p = 0.21, paired t-test, **Figure 1G**) on trials with the earliest licks and latest licks. Similarly, Cspk responses exhibited only a slightly smaller amplitude on trials with the slowest lick rates as compared to trials with the fastest licking (p = 2.62 ×10^-7^, paired t-test, **Figure 1H**), and similar timing across these two trial types (time from cue onset on trials with highest lick rates: 187.90 ± 2.71 ms; lowest lick rates: 185.01 ± 2.69 ms, p = 0.38, paired t-test, **Figure 1H**). Consistent with previous work^4,7^, these results suggest that the cue-associated Cspk responses after training are not primarily related to licking.

In addition to licking, mice also exhibited other body movements that could modulate Cspk activity in lobule simplex. We therefore used a piezoelectric sensor^4^ to measure body movement, and segregated trials with the most and least movement (**Figure 1I**). This analysis revealed a similar amplitude (p = 5.38 × 10^-5^, paired t-test, **Figure 1J**), and an almost identical timing of Cspk responses on trials with the least and most movements (time from cue onset on trials with least movements: 197.5 ± 2.72 ms, most movements: 201.17 ± 2.49 ms; p = 0.25, paired t-test, **Figure 1J**).

Consistent with the previous results^4^, these data suggest that Cspk activity shifts from the time of licking and reward consumption to the reward-predictive auditory cue, and that the cue-related Cspk responses are not driven by motor signals due to licking or body movements in trained animals. Moreover, the Cspk response to the auditory cue is similar to previously measured responses to a learned visual cue^4^, suggesting that CFs can represent reward-predictive stimuli of different modalities.

### Stimulus-blocking prevents learned Cspk responses

To test whether the CFs exhibit other signatures of reinforcement learning signals^15–19^, we next performed a stimulus-blocking experiment. A key prediction of reinforcement learning is that the transfer of neural activity from US to CS cannot occur unless there is an initial neuronal response to the US^13,14,20^. This requirement has important consequences. In particular, it means that a new CS (CS2) cannot be learned if it carries the same reward prediction as a CS (CS1) that has already been learned. This so-called ‘stimulus blocking effect’ is a hallmark of reinforcement learning systems, as it serves to appropriately bind the CS and US^21^. Moreover, this property ensures that an additional stimulus that provides no new information is not learned.

To test how CFs respond to a CS that does not carry new reward-predictive information, we introduced a new CS (CS2) after mice had already learned the initial auditory CS (CS1), and the US response was extinguished. For CS2, we chose a visual cue that we have previously demonstrated can be learned by CFs in this behavioral paradigm^4^. We then trained animals where reward delivery was predicted by the co-presentation of CS1 and CS2 (**Figure 2A**). After a similar number of trials (mean = 1006 ± 43.3 trials) and training sessions (mean = 8.5 ± 0.2 sessions, n = 6 mice) required for CS1 learning, we measured the conditioned licking response and climbing fiber response to CS1 alone, CS1+CS2, and CS2 alone (**Figure 2B-G**).

**Figure 2.**
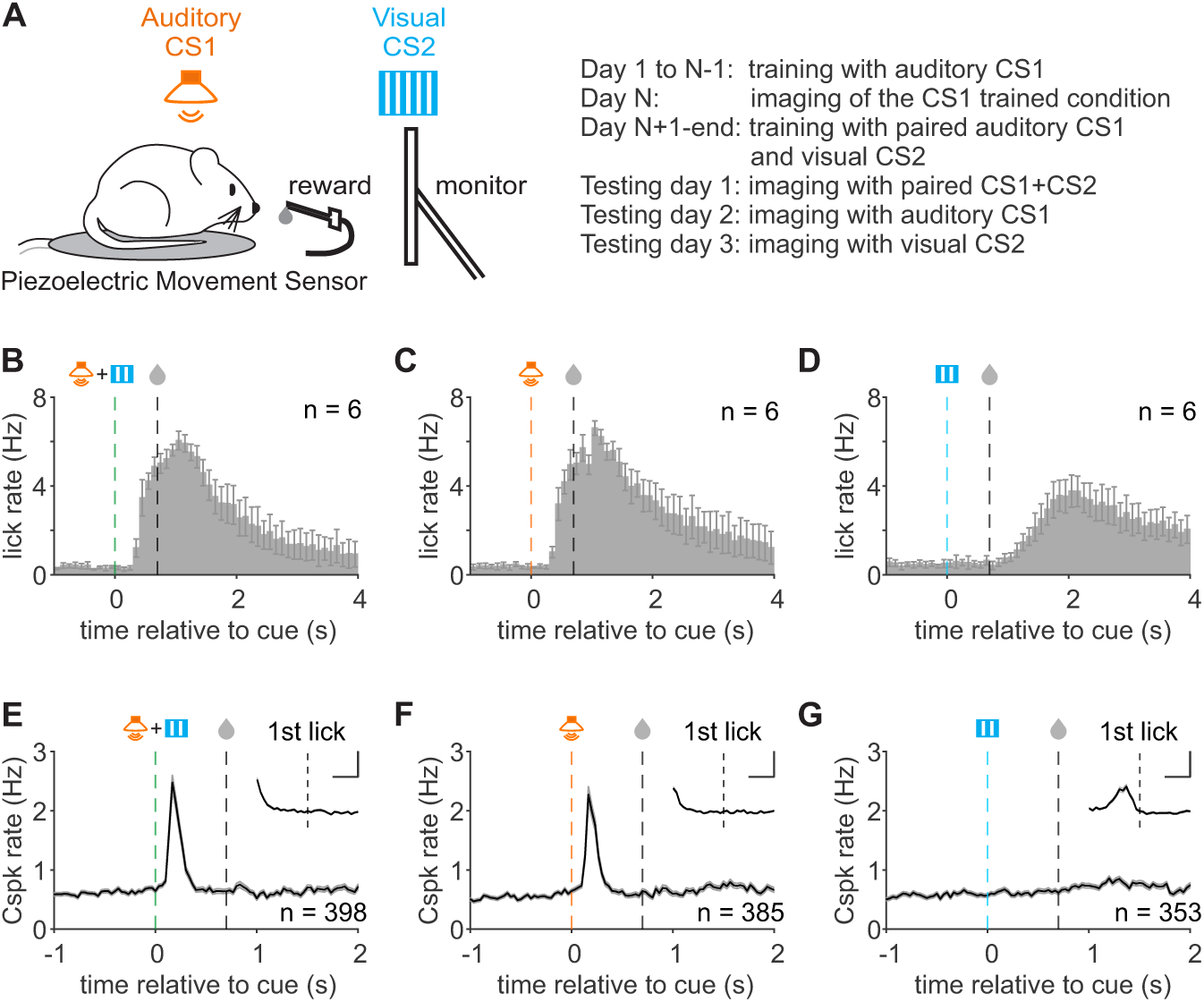
Cspk responses after training with a stimulus blocking paradigm. **A)** Task design (left) and training sequence (right) for mice initially trained on auditory CS1, and subsequently trained on the simultaneous presentation of auditory CS1 and visual CS2. Mice are then tested on sequential days with CS1+CS2, CS1 (auditory) alone and CS2 (visual) alone. **B)** Summary histogram of mean lick rates across mice aligned to cue onset in CS1+CS2 trained animals for CS1+CS2 presentation after learning (n=6 mice). Dashed lines represent the time of cue and reward respectively. **C)** Same as B) for CS1 presentation. **D)** Same as B) for CS2 presentation. **E)** PSTH of Cspk rate aligned to cue onset in CS1+CS2 trained animals for CS1+CS2 presentation after learning (n=398 dendrites). Inset, Cspk rate aligned to the first lick after cue presentation. **F)** Same as E) for CS1 presentation (n=385 dendrites). **G)** Same as E) for CS2 presentation (n=353 dendrites). All data are presented as mean +/- SEM.

We observed a predictive licking response to both CS1+CS2 and CS1 alone (CS1+CS2 reaction time 465.1 ± 29.3 ms, 86.0 ± 2.7% of trials with a predictive response, 8.9 ± 2.9% of trials with a reactive response; CS1 alone reaction time: 489.7 ± 31.2 ms, 84.3 ± 4.0% of trials with a predictive response, 10.0 ± 3.5% of trials with a reactive response; n = 6 mice; **Figure 2B,C, See also Figure S2**). In contrast, mice licked only in response to the US when CS2 was presented alone (reaction time: 1511.1 ± 88.4 ms, 5.0 ± 1.7% of trials with a predictive response, 90.8 ± 3.2% of trials with a reactive response, **Figure 2D, See also Figure S2**). This result is consistent with previous stimulus blocking experiments related to midbrain dopamine circuits^13,14,20^ and indicates that the redundant CS2 was not learned.

Consistent with the robust association of CS1 with reward, we observed significant Cspk activity after the presentation of either CS1+CS2 (p = 7.87 × 10^-65^, paired t-test, n = 398 dendrites, 6 mice, **Figure 2E**) or CS1 alone (p = 1.84 × 10^-49^, paired t-test, n = 385 dendrites, 6 mice, **Figure 2F**), and no Cspk response to the first lick after reward (CS1+CS2, p = 0.68; CS1, p = 0.90; paired t-test, **Figure 2E,F**). Notably, the amplitude of the Cspk response to CS1 (2.61 ± 0.14 Hz) was not significantly different from the response to CS1+CS2 (2.73 ± 0.12 Hz, p = 0.45, paired t-test), suggesting there was not an additive effect when both cues were presented simultaneously. Additionally, the time to peak Cspk activity was the same for CS1+CS2 and CS1 presentations (CS1+CS2 209.38 ± 2.25 ms; CS1: 224.43 ± 2.31 ms; p = 0.52, unpaired t-test). Thus, the learned climbing fiber response was unchanged as long as CS1 was presented.

In contrast, we did not observe cue-driven Cspk activity when CS2 was presented alone (p = 0.21, paired t-test, n = 353 dendrites, 6 mice, **Figure 2G**). Instead, in this condition we observed a significant response to the first lick after reward delivery (p = 7.15 × 10^-34^, paired t-test, **Figure 2G**). These results mirror what has been demonstrated in similar stimulus-blocking experiments for VTA dopamine neurons, fulfilling one key prediction of reinforcement learning signals.

### Optical IO inhibition impairs Cspk transfer from US to CS

While stimulus blocking experiments provide a correlative link between Cspk activity to the US and learned Cspk activity to the CS, they do not causally link these neural activity patterns. Thus, to test whether US driven Cspk activity is causally linked to learned, CS-driven Cspk activity, we repeated behavioral training while optogenetically inhibiting neurons in the inferior olive (IO) which are the source of the CFs to the cerebellar cortex. For these experiments, we unilaterally expressed the soma-restricted inhibitory anion-fluxing channelrhodopsin GtACR2^22,23^ in the contralateral IO, and implanted an optical fiber over this region (**Figure 3A, See also Figure S3**)^24,25^. To verify the efficiency of this optogenetic strategy *in vivo*, we first performed single-photon calcium imaging of Purkinje cell dendrites in lobule simplex. During optical inhibition, we observed a marked reduction of activity during the illumination period (p = 2.35 × 10^-4^, n = 10 trials, 5 mice, paired t-test, **Figure 3B,C**). Upon light offset, we observed a transient rebound response, followed by a return to baseline levels.

**Figure 3.**
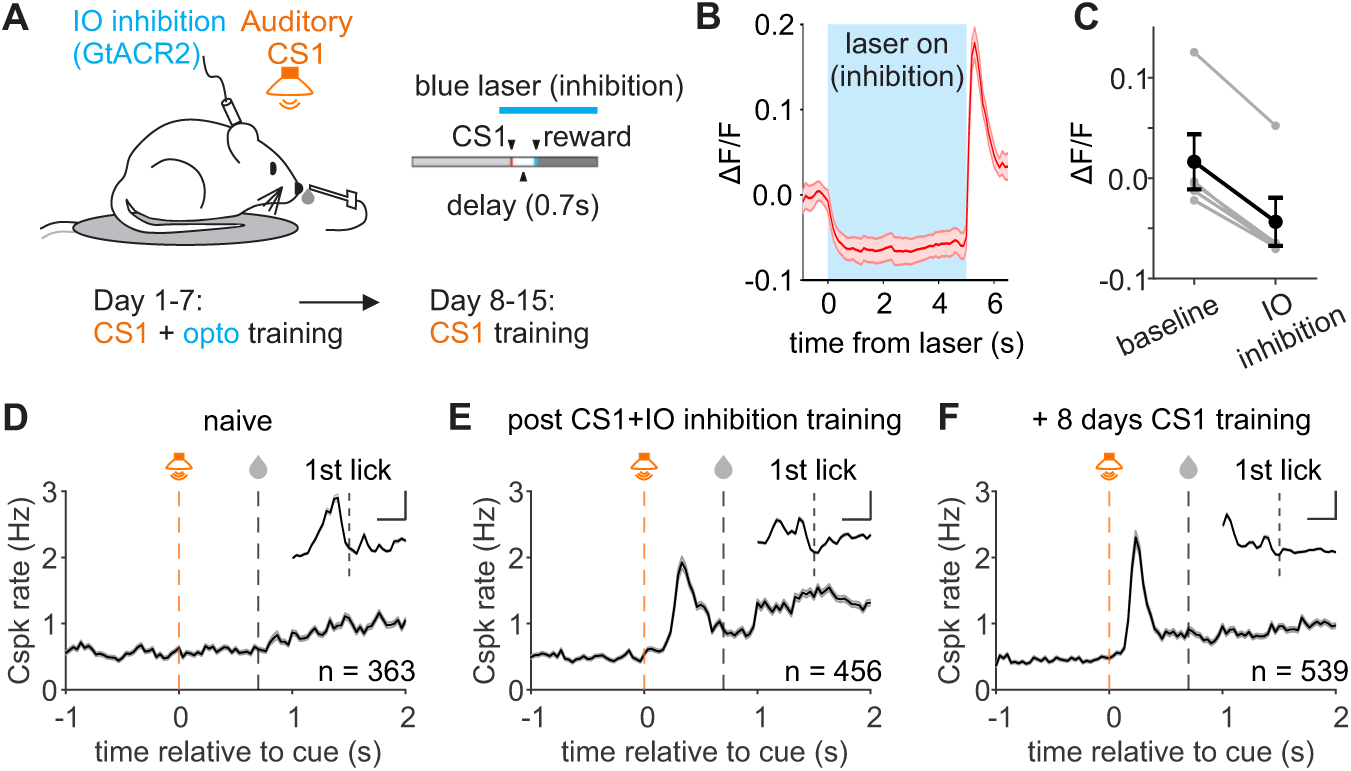
Cspk responses across training with optical inhibition of the contralateral IO. **A)** Task design for mice initially trained with GtACR2 inhibition of the IO during CS1 presentation, and subsequently trained with CS1 and no IO inhibition. **B)** Single photon imaging of bulk Cspk responses across mice (n=5 mice) during optical inhibition via fiber implant above the IO. **C)** Summary of mean Cspk response amplitude in individual mice (gray, n=5 mice) and across animals (black) in control and during optical IO inhibition. **D)** PSTH of Cspk rate aligned to cue onset for naive mice prior to GtACR2 training. Inset, Cspk rate aligned to the first lick after cue presentation (n=363 dendrites). **E)** Same as D) but in the same mice after training with IO inhibition (n=456 dendrites) **G)** Same as E) but in the same mice after additional training with CS1 (and no IO inhibition) (n=539 dendrites). All data are presented as mean +/- SEM.

In mice expressing GtACR2 in the IO, we first measured the Cspk response to the auditory CS in naive mice (**Figure 3D**). As previously, there was no cue-related Cspk response (n = 363 dendrites, 5 mice, p = 0.34, paired t-test, Figure 3c), but a significant US-related response (p = 4.75 × 10^-77^, paired t-test, **Figure 3D**). These results suggest that GtACR2 expression did not affect normal climbing fiber responses to reward.

Next, we trained animals while optically inhibiting the IO on each trial, beginning 400 ms before cue onset until 4 s after reward delivery, placing rebound activity well outside the trial structure (**Figure 3A,B**). Notably, we did not observe a significant lick response to the onset or offset of optical inhibition (onset p=0.42; offset, p = 0.11, paired t-tests, n = 34 training sessions, 5 mice, **Figure S3**).

After training during IO suppression (mean = 8 ± 0.5 sessions, n = 5 mice), we measured the Cspk responses in the absence of optogenetic inhibition in the following session. In stark contrast with normal training, we observed Cspk activity to both the auditory cue (p = 1.57 × 10^-79^, n = 456 dendrites, 5 mice, paired t-test, **Figure 3E**) and the first lick after reward delivery (p = 4.24 × 10^-81^, paired t-test, **Figure 3E**). Moreover, the cue-associated Cspk response occurred at a significantly longer latency than in normally trained animals (GtACR2 Cspk latency = 384.72 ± 2.31 ms; control Cspk latency = 181.03 ± 1.98 ms; p = 2.14 × 10^-243^, unpaired t-test). This intermediate response suggests an incomplete shift of Cspk activity from the US to CS. Thus, we continued training without optical inhibition to test whether normal training would generate responses similar to those in control animals.

Following 8 additional days of normal training, we observed Cspk activity in response to the auditory cue (p = 3.26 × 10^-96^, paired t-test, n = 539 dendrites, 5 mice, **Figure 3F**) that was significantly larger than it had been immediately after training with optogenetic inhibition (Post opto training: 2.45 ± 0.08 Hz; post additional training, 2.72 ± 0.09 Hz, p = 0.03; unpaired t-test, **Figure 3F**). Moreover, peak Cspk activity occurred earlier, in closer proximity to the auditory cue (280.58 ± 2.02 ms; p = 2.21 × 10^-169^, paired t-test), and the US-associated Cspk response was largely abolished (vs post optical training, p = 6.55 × 10^-14^, unpaired t-test, **Figure 3F**). Together, these results suggest that Cspk activity in response to the US contributes to the timing of the learned Cspk activity in response to a CS and the abolition of the Cspk activity in response to the US.

### Climbing fiber responses are consistent with variable reward expectations across trials

On average, Cspk activity is consistent with learned reward expectation in this task. However, reward expectation can also vary across trials, affording another means to test the relationship between behavior and neural activity. We therefore examined how Cspk activity varies across trials, leveraging both the responses immediately after optogenetic training and those after subsequent normal training. To do so, we segregated trials into predictive (pre-reward) and reactive (post-reward) licking.

Following optogenetic training, on trials with predictive licking nearly all individual PCs (93%) had Cspks in response to the cue (n = 423 dendrites; **Figure 4A**). Surprisingly, these cells also had a significant Cspk response prior to the first lick on these trials (p = 5.76 × 10^-96^, paired t-test, n = 423 dendrites, 5 mice, **Figure 4B**). However, the timing of this response was less closely associated with lick onset than in naive animals (from time of peak Cspk response to lick onset in trials with predictive licks in GtACR2 trained mice: 294.17 ± 2.27 ms; trials with reactive licks in naive mice: 115.06 ± 2.40 ms; p = 3.91 × 10^-315^, unpaired, one-tailed t-test, **Figure 4B** inset). Moreover, the cue-aligned Cspk response was also less closely associated with the auditory cue than it was in the same animals with additional normal training (from time of cue to peak Cspk cue response in GtACR2 trained mice: 436.56 ± 2.43 ms; in additionally trained mice: 316.03 ± 2.12 ms; p = 5.14 × 10^-188^, **Figure 4A** vs **4E**). This intermediate response suggests that most PCs show an incomplete / partial transfer of Cspk responses from the time of reward to the reward-predictive cue. In contrast, PCs that did not have Cspk responses to the auditory cue also did not show responses to the first lick after reward on predictive trials (p = 0.19, paired t-test, n = 33 dendrites, **Figure 4A,B**, grey). Thus, on predictive lick trials, the only response was intermediate, neither tightly linked to the auditory cue or reward delivery.

**Figure 4.**
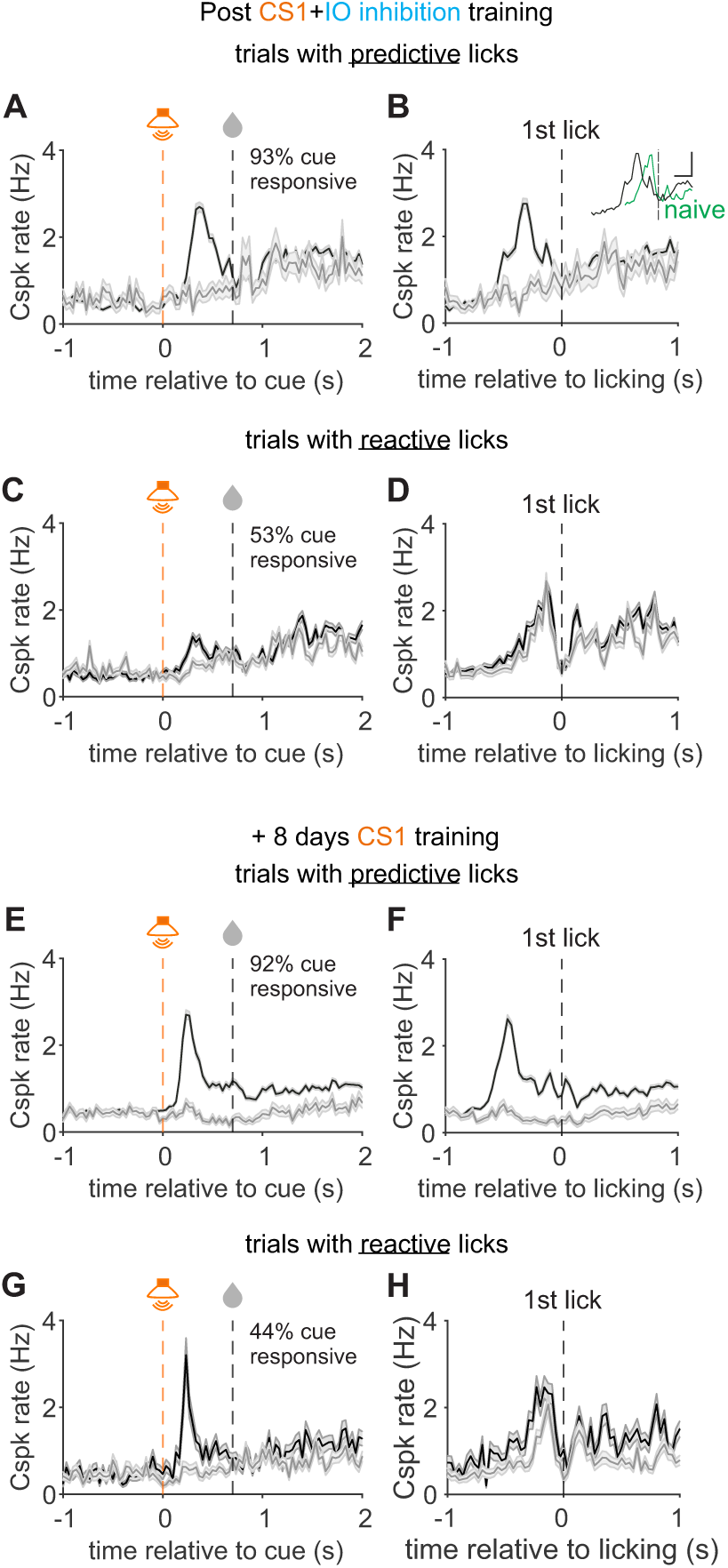
CF responses are consistent with reward expectation across trials. **A)** PSTH of Cspk rates on trials with predictive licking for PC dendrites that were cue responsive (black, n=423 dendrites) and non-cue responsive (grey, n=33 dendrites) aligned to cue onset in animals immediately after CS1 training with GtACR2 inhibition of the IO (n=5 mice). **B)** Same as A), but aligned to lick onset. Inset compares the same lick-aligned Cspks (black) to naive Cspk responses (green, from Figure 3D) **C)** PSTH of Cspk rates on trials with reactive licking for PC dendrites that were cue responsive (black, n=242 dendrites) and non-cue responsive (grey, n=214 dendrites) aligned to cue onset in animals immediately after CS1 training with GtACR2 inhibition of the IO. **D)** Same as C), but aligned to lick onset. **E)** PSTH of Cspk rates on trials with predictive licking for PC dendrites that were cue responsive (black, n= 497) and non-cue responsive (grey, n= 42 dendrites) aligned to cue onset in animals immediately after additional CS1 training (no IO inhibition, n=5 mice). **F)** same as E), but aligned to lick onset. **G)** PSTH of Cspk rates on trials with reactive licking for PC dendrites that were cue responsive (black, n= 237 dendrites) and non-cue responsive (grey, n=302 dendrites) aligned to cue onset in animals immediately after additional CS1 training (no IO inhibition, n=5 mice). **H)** same as G), but aligned to lick onset. All data are presented as mean +/- SEM.

Cspk responses were quite different on trials with reactive licking. First, far fewer PCs (53%) had a significant Cspk response to the cue (n = 242 dendrites, **Figure 4C**). Moreover, the amplitude of the Cspk response in those PCs was significantly smaller as compared to trials with predictive licking (p = 1.71 × 10^-35^, unpaired t-test, **Figure 4A** vs **4C**). However, both cue responsive and non-cue responsive PCs showed activity to the first lick after reward on reactive trials (cue-responsive dendrites: p = 2.09 × 10^-49^, n = 242 dendrites; non-cue-responsive dendrites: p = 1.02 × 10^-33^, n = 214 dendrites; paired t-tests, **Figure 4D**). This Cspk activity was more closely associated with lick onset as compared with trials with predictive licking, indicating that it was associated with anticipation or licking (from time of peak Cspk activity to lick onset in trials with reactive licks: 116.74 ± 2.19 ms; with predictive licks: 294.17 ± 2.27 ms; p = 9.95 × 10^-294^, unpaired t-test, **Figure 4B** vs **4D**). Thus, while most CFs showed a strong cue response on trials with predictive licking, they did not respond robustly as to the cue on trials with reactive licking, but instead responded as if the reward was unexpected, showing robust lick-associated responses. In this way, Cspk activity is consistent with tracking reward expectation across trials. Notably, Cspk responsivity to different task parameters (cue and reward) was spatially intermingled, and thus does not appear to be segregated into a clear microzonal organization.

To determine whether these across-trial relationships between reward expectation and Cspk activity were influenced by optogenetic manipulation of CFs during training, we compared responses to those observed following additional normal training. In this condition, most dendrites (92%) continued to have significant cue responses on trials with predictive licking (n = 497 dendrites; **Figure 4E**). However, when aligned to the first lick after reward this Cspk response was significantly further away from the lick, indicating that it was now fully associated with cue presentation, and no longer intermediate between cue and reward times (from time of peak Cspk activity to lick onset in trials with predictive licks in mice with additional training: 467.78 ± 2.02 ms; p = 1.77 × 10^-246^, unpaired, one-tailed t-test, **Figure 4F vs 4B**). Additionally, on trials with reactive licking a much larger response had developed to the cue (p = 1.61 × 10^-5^, unpaired, one-tailed t-test, **Figure 4G vs 4C**). Thus, the learned cue response was considerably more prominent on both trial types after normal training.

Together, these data further reveal how optogenetic inhibition impairs the development of learned Cspk responses. Moreover, they demonstrate that 1) PCs preferentially receive CF input on trials where licking will be predictive, and 2) CFs only respond to licking when it is reactive, again suggesting that these responses may be related to expectation. Thus, Cspk responses appear to track expectation across trials, in addition to across learning.

### Reward-associated Cspk activity influences learning

Our data suggest the Cspk activity can reflect reward expectation, but do these reward-related responses influence behavior or learning? To test this, we quantified lick responses immediately after optogenetic training, again after subsequent normal training in the same animals, and in separate animals with no optogenetic manipulation (**Figure 5**). Following optical inhibition of IO neurons across training, we observed that licking was reactive on many trials, and frequently did not peak until well after reward delivery (**Fig 5A-B, See also Figure S4**). In contrast, licking was predictive on the majority of trials following subsequent normal training, peaking before reward delivery (**Figure 5C, See also Figure S4**).

**Figure 5.**
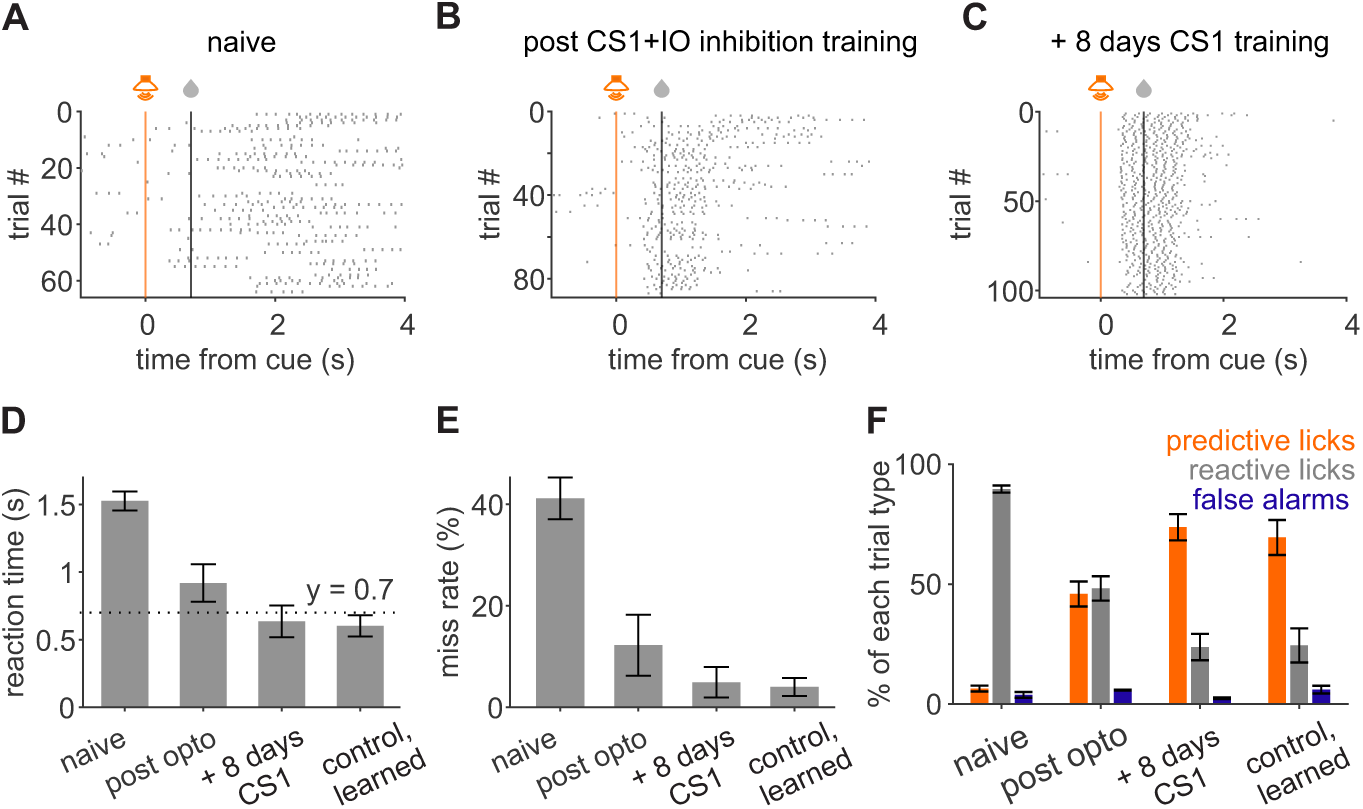
Training with IO inhibition impairs learned lick timing. **A)** Example lick raster, aligned to cue onset, for a mouse prior to CS1 training. **B)** Same as A, for the same mouse after CS1 training with GtACR2 inhibition of the IO. **C)** Same as A), for the same mouse after additional training with normal CS1 training (no IO inhibition). **D)** Summary of mean reaction times for the same mice before (naive), after GtACR2 training (post opto), and after additional CS1 training (+8 days CS1) (n=5 mice), as well as in a separate cohort that underwent only normal training (control, learned, n=6 mice). Dashed line is the time of reward. **E)** Same as D), for miss rates. **F)** Summary of the fraction of trials across mice with predictive licks (orange), reactive licks (gray), or early licks (false alarms, blue) during the trial for the same groups as in D). All data are presented as mean +/- SEM.

On average, following optogenetic training, mean reaction times occurred after reward delivery, favoring reactive licking (p = 0.09, n = 5 mice, Student’s one-tailed t-test, **Figure 5D, See also Figure S5**). These reaction times were significantly longer as compared with the same animals following subsequent normal training (p = 0.027, paired t-test, one-tailed, n = 5 mice) and as compared with animals trained without optogenetic inhibition (p = 0.007, unpaired t-test, one-tailed, n = 6 mice). Notably, learning had plateaued during the last three days of optical inhibition (reaction time p = 0.53, one-way ANOVA, n = 6 mice), suggesting that learning may not only be slowed, but that mice may not be able to achieve precisely timed behavior if reward-related climbing fiber input is suppressed (**Figure S5**).

There was also a trend toward higher miss rates following optogenetic training, though these were not significantly different from subsequent normal training or controls (optotrained vs subsequent normal training, p=0.14, paired t-test, one-tailed; vs control; p = 0.09, unpaired t-test, one-tailed Figure **5E**) In contrast, the percentage of trials with predictive licking (46 ± 12%) was dramatically lower than in the same animals with subsequent normal training (74 ± 12%, p=0.003, paired t-test, one-tailed, **Figure 5F**), or the control animals (69 ± 7%; p = 0.055, unpaired t-test, one-tailed, **Figure 5F**). Conversely, the percentage of trials with reactive licking following optogenetic training (48 ± 11%) was markedly higher than either the same animals with subsequent normal training (24 ± 12%, p=0.007, paired t-test, one-tailed, **Figure 5F**) or control animals (24 ± 7%, p = 0.050, unpaired t-test, one-tailed, **Figure 5F**). The large change in the number of predictive responses without a statistically significant change in miss rates suggests an impaired learning of predictive timing, but not an impairment in the ability to initiate licks as a result of CF inhibition during training.

Consistent with the observation that optical IO inhibition did not induce licking (**Figure S3**), we also did not find evidence that it impaired licking, as there was no difference in reaction times or rates of predictive licks from the last day of optical inhibition to the next day with no optical inhibition (**Figure S5**). This suggests that reward-related CF signals are primarily involved in learning, rather than the acute generation of predictive licks. Together, these results support a causal role for the climbing fiber response at the time of the reward in the learning of a well-timed conditioned licking response to the predictive cue.

## Discussion

Our data reveal multiple features of cerebellar climbing fiber signaling that suggest a broad role in reward processing and reward-based learning. First, we find that cerebellar CFs can generate learned, reward-predictive responses to auditory stimuli in this task, extending previous findings by indicating that CFs can learn sensory predictions for additional modalities. Second, we find that these learned reward predictions exhibit a key property in common with reward prediction error (rPE) signals in the VTA, in that they cannot generate learned responses to a redundant sensory cue. This so called ’stimulus-blocking effect’ has important consequences for learning, and suggests that the learned CF signals described here are not simply related to the salience of the cue, or its temporal association with reward. Third, our data also suggest a role for CF responses to unexpected reward in generating learned responses to reward-predictive cues, as optogenetically reducing CF activity impairs the transition from the US to the CS. And finally, our data also reveal a role for reward-related CF signals in learning and behavior, as optogenetic suppression of CF reward signals impairs the learning of predictively timed licking.

Previous work showed that CF responses can shift from the US to a visual CS^4^, and that CFs can respond to visual^7^ and auditory^5^ stimuli in mice trained on reward-based operant tasks. Notably, the learned CF response to an auditory cue in this study occurred with shorter latency and a larger amplitude as compared to previously measured visual responses^4^. We speculate that this may result from either the longer training period in this study or an enhanced salience or faster transduction time of the auditory stimulus as compared to the visual stimulus.

As with previous results, the US and CS-related CF responses could be disambiguated from neural activity associated with licking or body movement^4,7^. Thus, these data further support the conclusion that CF responses are related to the US in the naive condition and the CS in the learned condition. However, our current results also extend previous findings by explicitly testing whether these CF responses could simply be due to salience or simple a reward-related association. Reinforcement learning requires the detection of a discrepancy between the expectation and receipt of reward^13–15,21^. This attribute enables the learning of reward-predictive stimuli, but also prevents the learning of any stimulus that does not provide novel information about reward. Accordingly, simple pairing between stimulus and a reward is not sufficient for learning to occur ^3,14,20^. By demonstrating that a visual stimulus that is redundant with a previously learned, reward-predictive auditory stimulus cannot be learned, our data are in agreement with what has been shown in other reinforcement learning systems ^14,20^. Further, these data rule out the possibility that CF responses result only from the repeated exposure to a stimulus or its temporal proximity to reward.

Importantly, there remain other properties of reinforcement learning signals that have yet to be tested in the cerebellum. Indeed, even for midbrain dopamine neurons, substantial debate continues about the complex features of dopamine signaling, and how to reconcile the varied responses of dopamine neurons with the formal predictions of modern reinforcement learning theories such as temporal-difference learning^9,26^. Thus, while we conclude that at least some important features of CF activity are well-aligned with the predictions of reinforcement learning, much work remains to determine whether, when and how the cerebellum might participate in reinforcement learning.

Beyond the debate surrounding the computational properties of these CF responses, the circuit mechanism(s) that generate reward-driven and reward predictive CF activity also remain unclear. Broadly, CF reward-related responses could either be computed locally in the cerebellum or inherited from upstream brain regions. In this study, our optogenetic experiments indicate a contribution of cerebellar-olivary circuits in generating learned, reward-predictive CF responses. By inhibiting CF activity to unexpected reward, we observed an impaired transfer of CF signaling from the US to the CS. Notably, this manipulation did not block the generation of a CS response, but rather reduced its amplitude and temporal specificity, while at the same time allowing the US response to remain intact. This result may be due to incomplete block of CF activity, a requirement for more widespread circuit activity in upstream regions, or redundancy in the circuits that enable learned CF responses.

Indeed, previous studies have shown that the inferior olive receives descending inputs from multiple regions of the neocortex^27–29^. There is also evidence of polysynaptic input to the olive from the output of the basal ganglia^30^. However, there remains no clear evidence of projections from the VTA, despite evidence that the cerebellum may projects directly to the VTA^31^ and SNc^32^. Thus, current evidence suggests that the cerebellum may either act in parallel with midbrain dopamine neurons or contribute to their reward-predictive responses, but may not directly inherit reward-predictions from them.

The IO also receives inputs from the cerebellum that could contribute to CF reward computations. For example, excitatory inputs from the mesodiencephalic junction (MDJ) of the midbrain ^33^ receive signals from the deep cerebellar nuclei^34^, and also may integrate these with input from cortical pyramidal tract neurons ^29^. The IO also receives inhibitory inputs from the deep cerebellar nuclei, and has interneurons that may facilitate local computations^33,35^. Indeed, the inhibitory projection from the cerebellar nuclei to the IO has been shown to be necessary for suppressing behavioral responses to a redundant CS in eyelid conditioning^36^. Thus, it is possible that this projection may play a role in suppressing reward responses once they are expected. However, it remains unclear if this pathway might play any role in learned CS responses, and future work will be necessary to test the circuit pathways and mechanisms responsible for such responses.

Our optogenetic experiments also suggest a key role for reward-driven CF activity in generating well timed motor output. Specifically, we find that inhibiting the IO during learning impairs the ability to generate predictively timed licking in our task. These results are consistent with recent evidence linking the lateral cerebellum with learned, predictive licking^37–40^, and extend these findings by indicating a role for CF reward-related activity in learning. Here, we found that the effect of IO inhibition was specific to the learning of predictively timed responses, and did not affect lick rates outside of the task window. Moreover, by comparing the last day of IO inhibition with the first day of subsequent normal training, we found that IO inhibition did not directly impact lick reaction times or the fraction of trials with predictive licking. These same animals then proceeded to learn normally in absence of IO inhibition. Therefore, while reward-related CF signals correlate with expectation and facilitate learning, they do not appear to be causally involved in acutely generating the predictive licking behavior^37^. Notably, this interpretation does not rule out a contribution of CF activity to the precise timing of predictive licks^41^, and future work using high time resolution electrophysiology measurements on a trial-by-trial basis would be necessary to address this question. Together we conclude that reward-related CF signals can act as teaching signals for learning, as has been extensively demonstrated for traditional error driven CF signals in other tasks^1^. More broadly, these results are consistent with CF driven learning and cerebellar circuits contributing to predictive behaviors and anticipatory motor output^2^. Overall, our results suggest that cerebellar CFs can exhibit several properties in common with the reinforcement learning signals that have been characterized in the dopamine system^13,20^, and can play an important role in predictive learning.

## Resource Availability

### Lead contact

Further information and requests for resources and reagents should be directed to the lead contact, Court Hull.

### Material availability

No new reagents were generated as a result of this study.

### Data and Code Availability

- Due to the large size of these datasets, data that support the findings of this study are available from the corresponding author upon request.
- Code that supports the findings of this study have been deposited and are publicly available at: https://github.com/Glickfeld-And-Hull-Laboratories/ImagingCode-Glickfeld-Hull
- Any additional information required to reanalyze the data reported in this paper is available from the lead contact upon request

## Acknowledgements

This work was supported by grants from the following NIH institutes: NINDS 5R01NS096289 (CH), NINDS R01NS112917 (CH), NINDS R01-NS128054 (CH). Funding was also provided by the Ruth K. Broad Foundation (SJ). The funders had no role in study design, data collection and analysis, decision to publish or preparation of the manuscript. We thank Drs. Stephen Lisberger, Lindsey Glickfeld, Fan Wang, and Josh Huang for input throughout the project, and Mary Anne Hughes, Ziye Xu, and Wenjuan Kong for technical support.

## Author Contributions

C.H. and S.J. designed experiments. S.J. conducted experiments and analyzed data. C.H. and S.J. wrote and edited the manuscript.

## STAR Methods

### Experimental Model and Subject Details

All experimental processes involving mice were approved by the Duke Institutional Animal Care and Use Committee. Mice were singly or group-housed (<= 4 per cage) on 12-hour light/dark cycles.

Experiments were performed using adult mice (> P65) randomly selected from both sexes (n=7 male, 4 female) using Tg(PCP2-Cre)3555Jdhu mice (Jackson Labs, 010536).

### Method Details

#### Surgical Procedures

All surgical procedures were performed with an aseptic technique. Mice were injected with dexamethasone (3 mg/kg, or 4 mg/ml SQ) three to 24 hours before surgery. Ketamine (50 mg/kg) and xylazine (5 mg/kg) were administered by intraperitoneal injection for initial anesthesia, and then isofluorane (1-3%) was used to maintain the mice being anesthetized during surgery. A heating pad (TC-1000 CWE Inc.) was used to keep the body temperature at 37° Celsius.

The surgical procedures for craniotomy and virus injection were similar as stated by Heffley *et al.* ^7^. Specifically, a headplate (HE Parmer) was mounted to the skull with C&B Metabond (Parkell), then a 3 mm craniotomy was prepared over lobule simplex of the right cerebellar hemisphere (∼ 2.8 mm posterior and 1.4 mm lateral to lambda). To express GCaMP in Purkinje cell dendrites of lobule simplex, a glass micropipette was filled with virus AAV1.CAG.Flex.GCaMP6f.WPRE.SV40 (UPenn vector core, titer = 1.2 × 10^13^) or AAV1.CAG.Flex.GCaMP7f.WPRE.SV40 (Addgene 104496, titer = 1.5 × 10^13^) diluted 1:4 – 1:20 in ACSF (titer for injection: 7.5 × 10^11^). In each targeted lobule, 150-200 nL virus was injected at 2-4 sites using the following depths: 500 µm, 380 µm, 250 µm, and 100 µm. The injection rate was 30-50 nL/min. For a subset of animals with a craniotomy over lobule simplex, the virus injection was performed at an approach angle of 40 degrees with the same injection rate, volume, and depths ^5^. A glass window consisting of two 3 mm and a 5 mm coverslip (Warner Instruments No. 1) was implanted using C&B Metabond.

To express an inhibitory opsin in the inferior olive, the skull was leveled, and the following coordinates were used for the burrhole: x = 0.4 mm × (distance between bregma and lambda/6.85 mm); y = 6.85 mm × (distance between bregma and lambda/6.85 mm). Virus AAV1.αCaMKII.stGtACR2-FusionRed (Addgene 105677-AAV1, titer = 1.2 × 10^13^) was injected into the brainstem at a depth of 5.8 mm and 5.4 mm with a rate of 40-60 nL/min. The injection volume for each depth was 300 nL. An optical fiber (CFMLC21U-20, 105 µm, 0.22 NA, 6 mm in length, Thorlabs; or R-FOC-BL100C-22NA, 100 µm, 0.22 NA, 6.5 mm in length, RWD Life Science Co) was then implanted using the same coordinates using C&B Metabond. Mice were then headposted with a headplate (HE Parmer). After seven to 10 days of recovery, a second surgery including craniotomy over lobule simplex and GCaMP injection was done.

After the surgery, all animals were post-operatively monitored by our lab personnel for 5 days, and cefazolin (50 mg/kg) and buprenex (0.05 mg/kg) were injected every 12 hours for 2 days. At least a full week of recovery period was allowed before further procedures.

#### Behavior

All mice were habituated to head restraint for at least 5 days (10 min – 1h/day) before training and imaging. Imaging mice were allowed for at least 13 days after cranial injections to allow sufficient expression levels of GCaMP. Training and imaging sessions are shorter than 90 minutes to keep the animals engaged.

Mice were water deprived for at least 3 days until body weight was reduced and stabilized at around 85% of the original weight. All behavioral training occurred during the light cycle. Animals were head-fixed facing a reward delivery tube and a monitor. To measure body movement, a piezoelectric sensor (C.B. Gitty, 41 mm ‘jumbo’ piezo) was placed beneath the animal, and the time course of voltages was collected through MATLAB (Mathworks).

During the stimulus-blocking experiment, mice were first trained to associate an auditory cue (5 kHz, 75 dB, 200 ms tone) with a saccharine reward (approximately 2.5 µl) that was delivered 777 ms after the cue onset. Behavioral parameters including the onset of cue presentation, time of reward delivery, and time of licks were collected through MWorks (http://mworks-project.org) and MATLAB (Mathworks). Training of the auditory cue consisted of 1089.3 ± 103.6 trials (n = 9 ± 0.7 sessions, n = 6 mice), with an inter-trial interval (ITI) of 12 – 30 s. Learning of the association between the tone and the reward was defined by a stabilized reaction time of shorter than 650 ms since cue onset and miss rates below 15%. Once the mice learned the association, the same cohort of mice was then trained under a paired-stimuli condition. In this condition, the animals were presented with both a visual cue (600 ms, high contrast black and white vertical grating) and the same tone simultaneously during training, and were trained for a similar number of trials that took for the animals to learn the tone. Behavioral responses to the paired stimuli, auditory cue only, and visual cue only were then tested respectively and orderly in the imaging sessions following training. Training of the paired stimuli consisted of 1006 ± 43.3 trials (n = 8.5 ± 0.2 sessions, n = 6 mice).

During the training sessions with optogenetic suppression of the inferior olive, a 450 nm blue laser (Optoengine, MGL-III-450) was coupled to the optical fiber using a black ceramic sleeve (R-MS-1.25, 1.25 mm, black, RWD Life Science Co). The laser was turned on 400 ms before the cue onset, and terminated 4s after the reward delivery. A laser power of <1 mW was used. The black ceramic sleeve blocked the blue light and avoided the animals’ detection of the light delivery. The training with optogenetic suppression and auditory tone consisted of 742.2 ± 57.5 trials (n = 8 ± 0.5 sessions, n = 5 mice), and the training with only auditory tone consisted of 795.2 ± 64.9 trials (n = 8 ± 0.5 sessions).

#### Calcium Imaging

A customized microscope (Sutter SOM) and a CMOS camera (Qimaging, Rolera em-c^2^) were used for single-photon imaging. Excitation (470 nm) of the GCaMP6f or GCaMP7f was from an LED (ThorLabs, M470L3) through a 5x or 4x objective (Mitutoyo, 0.14NA), and the fluorescence signals were collected by a green filter (520 nm center wavelength, 25 mm diameter, 36 nm bandwidth, OD 6, Edmund Optics 67-030) with a 10 Hz frame rate. The field of view (FOV) was 3.5 × 3.5 mm (1002 × 1004 pixels).

In each trial, blue light was delivered to the inferior olive by a 450 nm blue laser (Optoengine, MGL-III-450) coupled to the optical fiber using a black ceramic sleeve (R-MS-1.25, 1.25 mm, black, RWD Life Science Co). To match the behavior paradigm, the laser was turned on for 5 s, and the ITI was 30 s. The imaging sessions consisted of 15 trials, and a laser power of <1.5 mW was used.

A resonant scanning microscope (Neurolabware) and a 16x objective (Nikon CFI75 LWD 16xW 0.80NA) were used to measure *in vivo* Ca^2+^ signals. A polymer (MakingCosmetics, 0.4% Carbomer 940) was used to preserve the immersion solution during two-photon imaging. A 920 nm excitation light from a Ti:Sapphire laser (Spectra Physics, Mai Tai eHP DeepSee) was raster scanned onto the brain at a 30 Hz frame rate via a resonant galvanometer (8 kHz, Cambridge Technology). Emitted photons were collected through a green filter (510 ± 42 nm band filter (Semrock)) onto GaAsP photomultipliers (H10770B-40, Hamamatsu). The microscope was controlled using Scanbox software (Neurolabware), and the FOV was 1030 μm x 581 μm (796 x 264 pixels). The laser power onto the brain was < 40 mW when imaging Purkinje cell dendrites.

#### Histology

Animals were administered with a cocktail of ketamine/xylazine (200mg/kg & 30mg/kg, respectively, IP) and at least 15 minutes were allowed until breath rate was < 60 bpm. Perfusion was then done transcardially with 10 - 15 ml 1x PBS followed by 4% paraformaldehyde (PFA) in 1x PBS. Immediately after the perfusion, dissection was done to isolate the brain, and the same PFA solution was used for post-fixation for 12 – 48 hrs (4°C). 100 μm sagittal slices were prepared using a vibratome (Pelco 102). For sections that included both the cerebellum and the brain stem, the brain was secured in a 2% agar block during slicing to maintain the section intact. Slices were first stained with DAPI (EMD Millipore Corp, DAPI, Dihydrochloride, 268298-10 mg) and then mounted with mounting medium (Southern Biotech Fluoromount-G) or directly mounted with mounting medium (Southern Biotech DAPI Fluoromount-G). Slides were imaged using a fluorescence (Nikon Eclipse 80i) or confocal microscope (Leica SP8).

### Quantification and Statistical Analysis

Data were analyzed and visualized using MATLAB (Mathworks) code. Unless stated otherwise, data were presented as mean ± S.E.M, and differences were considered statistically significant when p<0.05. Data distribution was assumed to be normal but was not formally tested. Analysis code was uniformly applied across experimental conditions.

#### Behavior

Licks were counted in an ‘all-or-none’ manner using a custom lick sensor ^4,7^, and all lick times during each trial were extracted. Lick bursts were defined as more than 3 licks with a 500 ms window. To determine whether the animals learned the task, licks were aligned to cue onset, and when calculating the following parameters, the time of the cue onset was considered 0 ms. The reaction time was defined as the onset time of the initial lick burst (or initial lick if no lick bursts were observed in a trial), and was calculated by averaging trials with a first lick or lick burst that occurred 200 ms - 3 s. Predictive licks (conditioned licking response before reward) were defined as the first lick or lick burst between 200 ms and 700 ms. Trials with reactive licks (after reward delivery) were defined as the first lick or lick burst later than 700 ms. Trials with unconditioned licking response were defined as the first lick after 700 ms and before 1700 ms. Early licks (false alarms) were defined as licks <200 ms from the cue. The miss rate was calculated by dividing the number of trials without a lick within 1.7 s by the total number of trials. For the cohort of mice that were trained with IO photoinhibition, licks were also aligned to the laser offset to determine whether the licking was perturbed by the optogenetic stimulation or the rebound climbing fiber activity after laser offset. Lick rates were measured as the mean lick rates during 1.5 s immediately before and after the cessation of illumination, and was compared across the 2 windows.

Trials were segregated according to lick responses in two different ways. First, to segregate trials according to the lick response latency, times of the initial lick burst (more than 3 licks within a 300 ms window) after the cue onset in each trial were identified, and then trials were divided into quartiles. Second, to segregate trials according to the lick frequency after reward delivery, the lick frequency of each trial was first calculated by dividing the total number of licks by the total length of the lick response window (from 100 ms to 2.5 s after reward delivery), and then trials were segregated into quartiles. To segregate trials according to body movements, mean voltage amplitude during the pre-reward window (from 100 ms to 800 ms after cue onset) was first calculated for each trial, and then trials were segregated into ten percentiles.

#### Single-photon mesoscale imaging

Lobule identity was identified based on craniotomy. Regions of interest (ROIs) were manually selected within each lobule according to the fluorescence of GCaMP signals. Normalized fluorescence (ΔF/F) was measured using the cumulative fluorescence within each ROI. For the optogenetics during single-photon mesoscale imaging, baseline fluorescence (F) was calculated by averaging the fluorescence across trials during the 1s before the laser onset. To be consistent with the trial structure during the behavioral sessions with optogenetic inhibition, the ΔF/F during laser on was measured as the mean of 1.1 – 5 s after laser onset.

#### Two-photon imaging

The extraction of individual Purkinje cell dendrites was performed as stated by Heffley *et al.* ^7^. Briefly, the following steps were taken: (1) Motion correction of the X-Y plane was done by registering image stacks to one reference image made by an averaged 5000-frame image. (2) Signals from individual dendrites were isolated by principal component analysis (PCA) and independent component analysis (ICA). (3) To further segment dendritic signals from the noise, a threshold of 3 - 3.5% was used when smoothing spatial filters from ICA. (4) Nearby pixels with a correlation coefficient > 0.8 were combined into a single dendrite to remove overlaps between dendrites.

To identify events used for the estimation of complex spike rates, deconvolution of the raw fluorescence was conducted on a cell-by-cell basis using MATLAB OASIS ^42^. The ‘FOOPSI’ method was used to account for the shape of the Ca^2+^ transients using GCaMP as a Ca^2+^ indicator. Before identifying complex spike events using deconvolution, Butterworth filter (high pass, order 3, cut off frequency 0.035 Hz) was applied to the raw fluorescence data to remove slow increase of fluorescence over time due to laser artifacts. A threshold of 3 standard deviations above the baseline was used across sessions for each experiment to account for the size of the events. Individual Purkinje cell dendrites were excluded based on the following criteria: (1) The fluorescence was above baseline consecutively for longer than 60 s. These dendrites typically exhibited a high level of fluorescence signals and a series of events that might not be related to climbing fiber activity. (2) The baseline fluorescence was below the level that allows the detection of any events. These cells were typically on a different z-plane or had a low level of GCaMP expression. F was calculated on a cell-by-cell basis, by averaging the fluorescence across trials during 50 frames (1.67s) before the cue onset. Spike rates were aligned to cue onsets or first lick after reward delivery for the mean peri-stimulus histograms (PSTHs).

To determine whether complex spike activity exhibited a significant response and whether the response latency and amplitude was significantly different across conditions, baseline activity of each cell was measured as the mean spike rates during 200 ms immediately before cue onset across trials. The response window for cue was defined as 100-300 ms after cue onset, and the response window for licking was defined as 200 ms immediately before the first lick after reward delivery. The response for each individual Purkinje cell dendrite was measured as the mean spike rates during the response window across trials, and the response latency for each individual dendrite was measured as the average time between the cue onset and the peak of Cspk rate during the response window across trials. The amplitude of Cspk response to cue was measured as the mean of peak Cspk rate across cells during the response window subtracted by the Cspk rate during baseline.

For the cohort of mice trained with optogenetic inhibition, to segregate Purkinje cell dendrites according to their responses to the cue, the baseline ΔF/F of each dendrite was measured by using the mean ΔF/F of 500 ms before the cue onset across trials, and the response amplitude was measured by using the peak ΔF/F during the window from 150 to 500 ms after the cue onset. Dendrites with a response amplitude of at least 3 standard deviations above the baseline were considered cue responsive. To determine whether complex spike activity exhibited a significant response and whether the response latency and amplitude was significantly different across conditions, baseline activity of each cell was measured as the mean spike rates during 300 ms immediately before cue onset across trials. Consistent with previous analysis, the response window for cue in the naïve condition was defined as 100-300 ms after cue onset. Since the time of the peak Cspk rate varies across conditions after training, the response window for cue in all other conditions was defined as 100 ms before and after the time/frame of peak mean Cspk rate across cells after the cue. When all Purkinje cell dendrites were considered, the response window for licking was defined as 200 ms immediately before the first lick after reward delivery. When comparing the time of peak Cspk rate before the first lick after reward of cue-responsive versus non-cue-responsive cells, the response window was defined as 100 ms before and after the time/frame of peak mean Cspk rate before licking.

---

**Figure S1.**
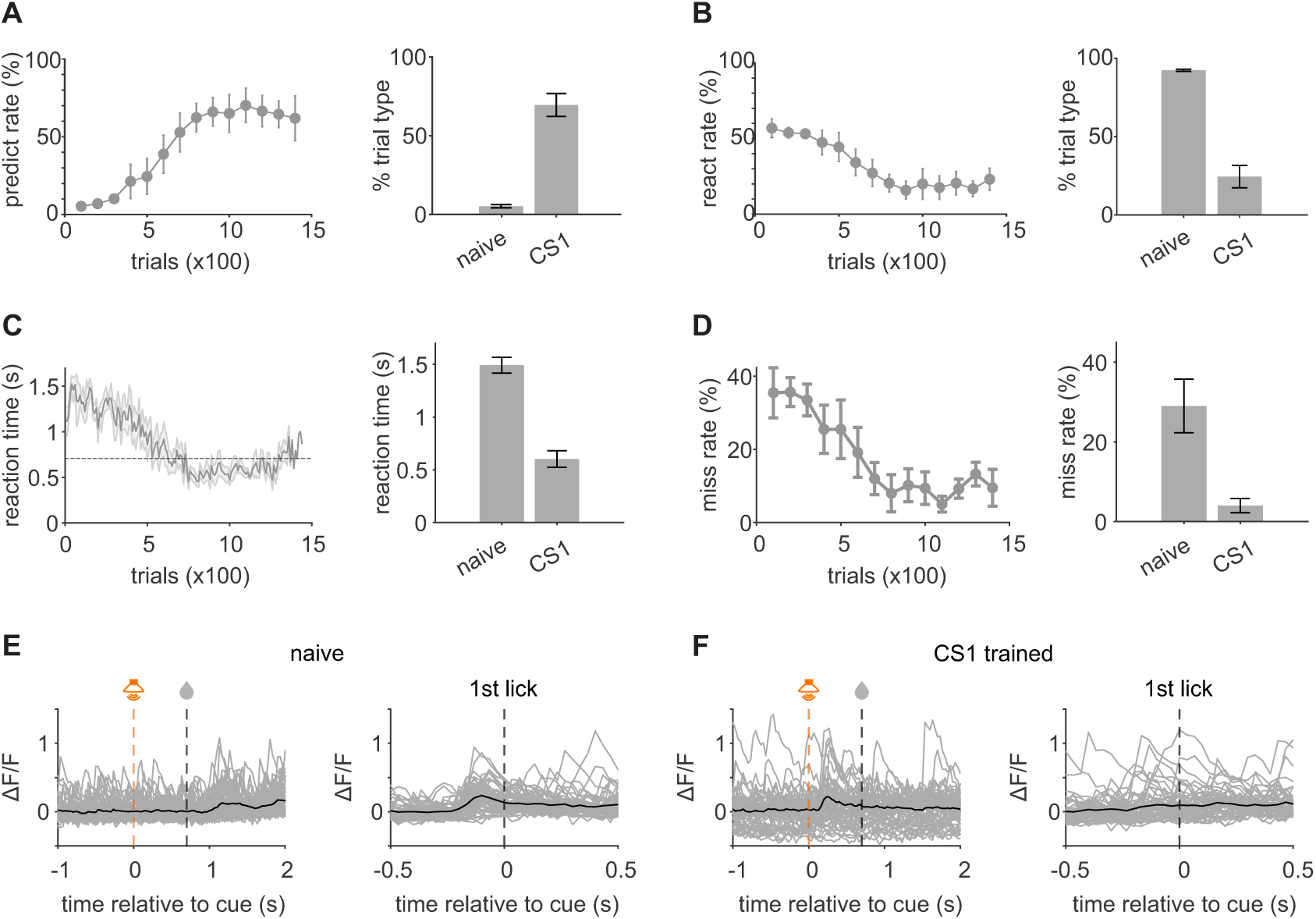
Summary of auditory CS1 behavioral training and example Cspk responses quantified by normalized GCaMP fluorescence (ΔF/F), related to Figure 1. A) *Left,* fraction of trials across mice with predictive licks (conditioned licking before reward), defined as the first lick or lick burst between 200 ms and 700 ms (n=6 mice). *Right*, summary of mean fraction of predictive licking before and after learning. B) *Left,* fraction of trials across mice with reactive licks (unconditioned licking after reward), defined as the first lick or lick burst after 700 ms. *Right*, summary of mean fraction of reactive licking before and after learning. C) *Left*, mean reaction time, defined as the first lick burst following the auditory cue, across animals. Dashed line is the time of reward. *Right*, summary of mean reaction times before and after learning. D) *Left*, mean miss rates, defined as the fraction of trials with no licks within 1.7 s after the cue, across animals. *Right*, Summary of mean miss rate before and after learning. **E)** Normalized GCaMP responses from an example segmented PC dendrite aligned to cue onset (left) and lick onset (right) for individual trials (gray, n = 100) and averaged across trials (black). **F)** Same as C), but for an example segmented PC dendrite (n=100 trials) in the same mouse after CS1 training. All data in A-D) are presented as mean +/- SEM.

**Figure S2.**
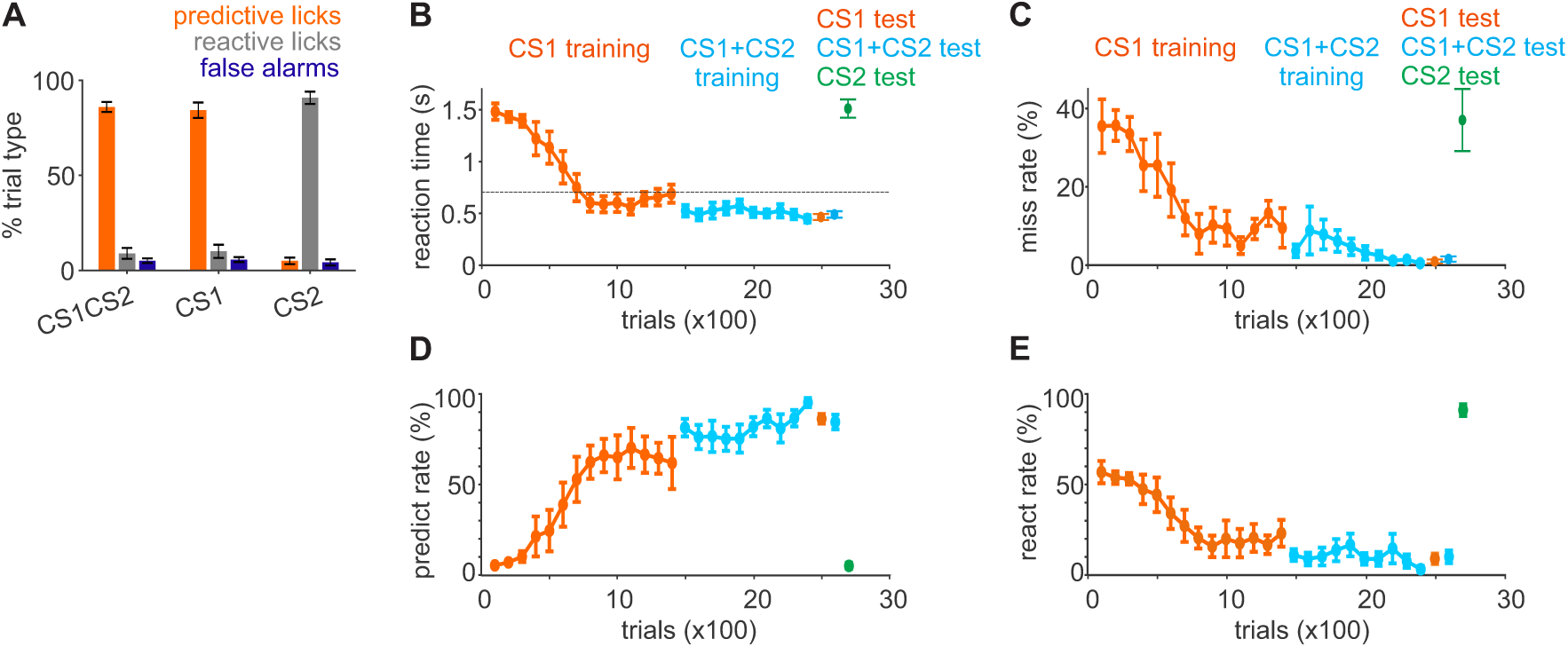
Summary of trial outcome and behavioral training for stimulus blocking paradigm, related to Figure 2. **A)** Summary of the fraction of trials across mice with predictive licks (orange), reactive licks (gray), or early licks (false alarms, blue) during the trial (n=6 mice). **B)** Mean reaction times across animals. orange=CS1 presentation, blue=CS1+CS2 presentation, green=CS2 presentation. Dashed line is time of reward. **C)** Same as B), for mean miss rates, defined as the fraction of trials with no licks within 1.7 s after the cue, across animals. **D)** Same as B), for mean predict rates across animals. **E)** Same as B), for mean react rates across animals. All data are presented as mean +/- SEM.

**Figure S3.**
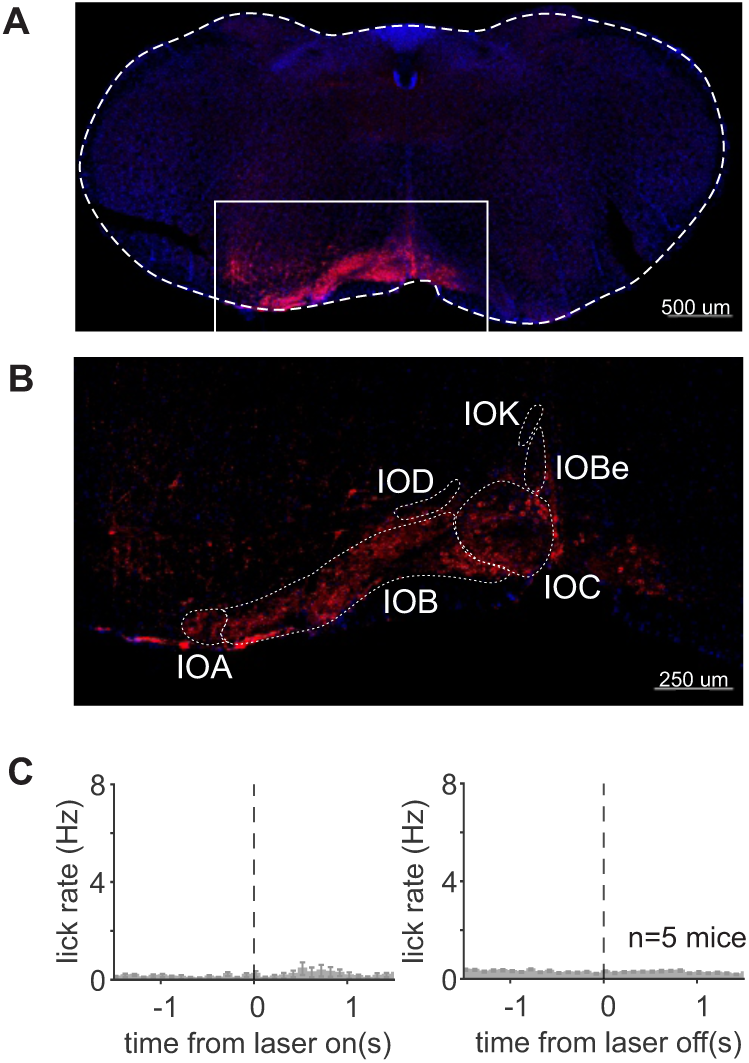
Expression of GtACR2 in in the IO, and effect of GtACR2 activation on licking, related to Figure 3. **A)** Coronal slice showing IO expression of fusion red (red) associated with GtACR2 expression following injection of pAAV-CKIIa-stGtACR2- FusionRed and DAPI staining (blue). Dotted line represents the outline of the brain section, and solid box represents the expanded field of view in B). **B)** Expanded field of view from A) showing expression in the inferior olive. Dotted lines represent IO subdivisions; IOA: subnucleus A of the medial nucleus; IOB: subnucleus B of medial nucleus; IOBe: beta nucleus of the medial nucleus; IOC: subnucleus C of medial nucleus; IOD: dorsal nucleus; IOK: cap of Kooy of the medial nucleus. **C)** *Left*, summary histogram of mean lick rates across mice expressing GtACR2 in the IO, aligned to laser onset (laser stim = 1.5 mW, 5 s, n=5 mice). *Right,* same as for C) but aligned to laser offset. Data are presented as mean +/- SEM.

**Figure S4.**
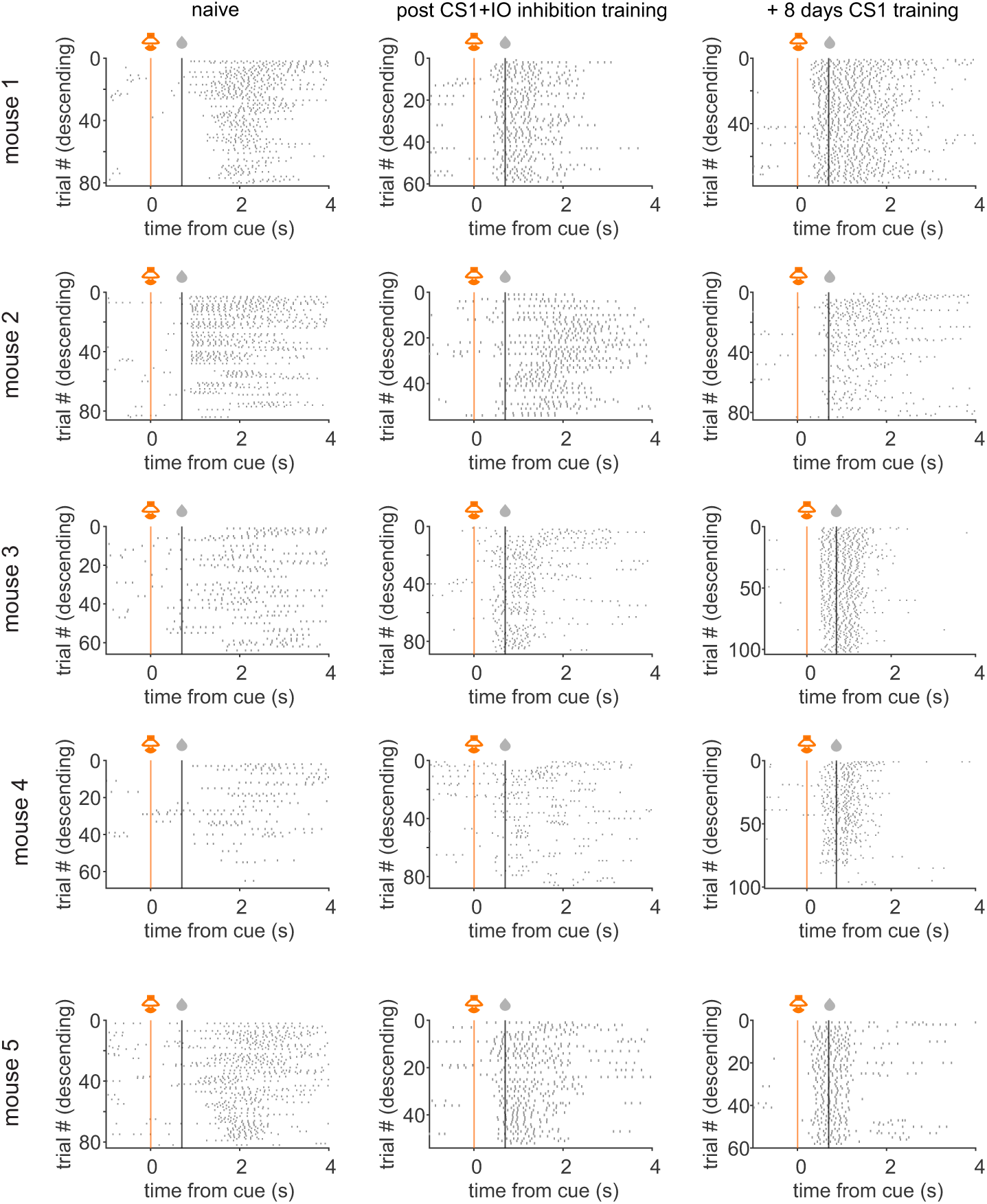
Lick rasters for all mice that underwent training with GtACR2 inhibition of the IO, related to Figure 5. Lick rasters, aligned to cue onset, for each mouse (individual rows) prior to training with GtACR2 inhibition of the IO (left), after CS1 training with GtACR2 inhibition of the IO (middle) and after additional CS1 training without GtACR2 inhibition of the IO (right).

**Figure S5.**
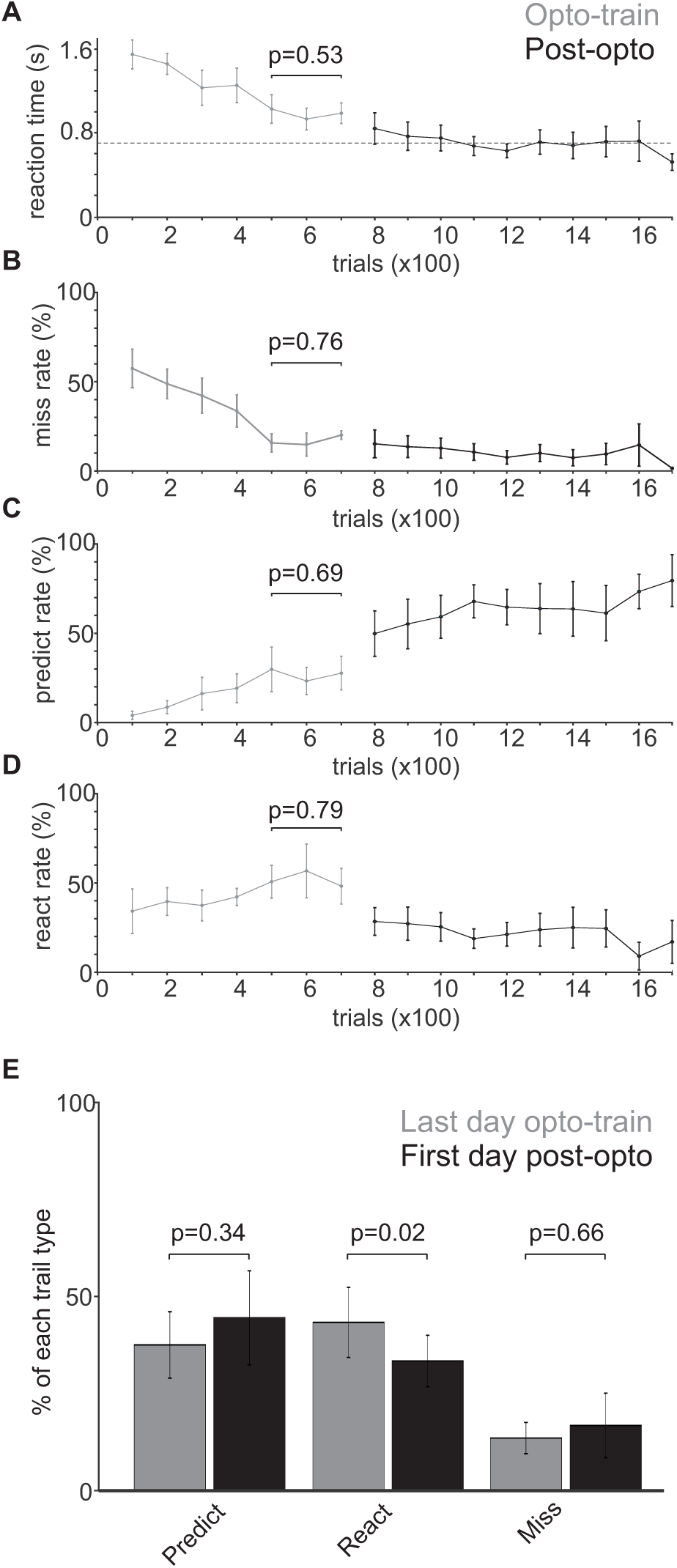
Summary of trial outcome across mice and sessions during GtACR2 training (gray) and subsequent normal training (black) (n=5 mice), related to Figure 5. **A)** Mean reaction times across training. Dashed line is time of reward. P-value is from one-way ANOVA on the last three sessions of GtACR2 training. **B)** Same as A, for miss rates. **C)** Same as A, for predictive lick rate. **D)** Same as A, for reactive lick rate. **E)** Summary of trial outcomes comparing the last day of GtACR2 training (gray) and the first day of subsequent normal training (black). P-values are from paired t-test. Data are presented as mean +/- SEM.

